# Label-free Detection and Characterization of Metabolism in Fresh and Cryopreserved Macrophages

**DOI:** 10.1101/2025.01.13.632815

**Authors:** Daniela De Hoyos Canales, Linghao Hu, Alex J. Walsh

**Affiliations:** Department of Biomedical Engineering, Texas A&M University, College Station, TX

**Keywords:** autofluorescence imaging, macrophages, metabolism, NADH, FAD

## Abstract

Cryopreservation is a widely used technique to preserve biological samples for extended periods of time at low temperatures. Even though it is known to have significant effects on cell viability, its effect on their metabolism remains unexplored. Studying how cryopreservation influences the metabolism of cells is important to guarantee the reliability of samples transported between sites for analysis. Optical metabolic imaging allows for the study of cellular metabolism in a label-free manner by using the autofluorescence properties of nicotinamide adenine dinucleotide (NADH) and flavin adenine dinucleotide (FAD), two metabolic coenzymes. The goal of this research is to study the metabolic changes in macrophages after cryopreservation and compare these results with freshly isolated macrophage samples to evaluate if the metabolic data is retained after cryopreservation. The metabolism of macrophages was analyzed with fluorescence lifetime imaging microscopy (FLIM) using a multiphoton microscope. Monocytes were isolated from human whole blood and separated into two groups. In Group 1, freshly isolated monocytes were differentiated into macrophages using macrophage-colony stimulating factor (M-CSF) over 8 days. In Group 2, isolated monocytes were cryopreserved after separation from whole blood and then thawed and stimulated with M-CSF for 8 days. Single-cell analysis showed there are significant changes in FLIM parameters between the groups suggesting that the metabolic data of macrophages is altered after a cryopreservation and cell thawing cycle. Our research bridges the current gap by studying the metabolic changes of cells after cryopreservation using a non-invasive and label-free imaging technique.

## 1. Introduction

Cryopreservation is a widely used method to store cell samples for extended periods of time. The cells are stored at very low temperatures, which allows the chemical processes of these cells to stop. This technique is commonly used by researchers to store cell samples for later use resulting in minimal damage to the cells [1]. While the impact of cryopreservation on the viability of cells has been studied, its effect on the metabolism of cells has not been explored [1]. This research study aims to bridge the current knowledge gap by using FLIM to analyze the metabolism of cells after a cryopreservation and thawing cycle. This study will help answer the questions of whether there is a change in metabolism of cells after cryopreservation by comparing data of fresh macrophages against that of cryopreserved macrophages.

The data collected in this study was gathered using FLIM to analyze the metabolic data of fresh and cryopreserved macrophages. Studying how cryopreservation influences cellular metabolism is important for several research applications, such as FLIM, where fresh and live cells samples are not always available. It is common for cell samples to be cryopreserved for long periods of time before being thawed and analyzed [2]. It is crucial to effectively analyze the metabolism of cells over time and after a cryopreservation cycle, as this can better studies that do not have access to fresh cells for analysis. It is also important to highlight that studying the metabolic state of cells is useful to gain information of disease conditions and therapeutic results [3].

The imaging of nicotinamide dinucleotide (NADH) and flavin adenine dinucleotide (FAD), two metabolic co-enzymes, provides a label-free method to study cellular metabolism. NADH and FAD are auto fluorescent with peak excitation and emission wavelengths around 340/550nm and 445/550nm, respectively. The fluorescence emissions of NADH and NAD(P)H, the phosphorylated form of NADH, are indistinguishable; therefore, the measured fluorescence signal of both molecules is generally denoted as “NAD(P)H” [4]. Similarly, the optical redox ratio (ORR) is commonly used to quantify the redox state of cells using NAD(P)H and FAD fluorescence intensities [3, 5].

## 2. Materials and methods

Monocytes were isolated from human whole blood and divided into two groups for this research study. The first group of monocytes, Group 1, were stimulated with the M-CSF cytokine over 8 days to allow for differentiation into macrophages. In Group 2, macrophages were cryopreserved after isolation from whole blood and then thawed for culture with cytokine over a period of 8 days.

### 2.1 Isolation of monocytes from human whole blood

Human whole blood was obtained from Stem Cell Technologies and processed to isolate fresh monocytes (Fig. 1). This separation was done using density gradient separation and the RosetteSep™ Human Monocyte Enrichment Cocktail from Stem Cell Technologies. To separate the monocytes from whole blood, 0.5 to 3 mL of blood were placed in a centrifuge tube along with EDTA to a concentration of 1 mM of EDTA. Similarly, 50 µL/mL of blood was added and mixed into the solution before incubating the tube for 10 minutes at room temperature. An equal volume of recommended medium (PBS containing 2% FBS and 1 mM EDTA) was added to tube. The sample is then transferred into a SepMate™ tube containing 4.5 mL of a density gradient separation medium, Lymphoprep™. The sample was centrifuged for 10 minutes at 1200 x g. The supernatant, containing the isolated monocytes, was transferred into a new tube to wash the monocytes with recommended medium by centrifuging the cells for 10 minutes at 300 x g.

**Figure 1.**
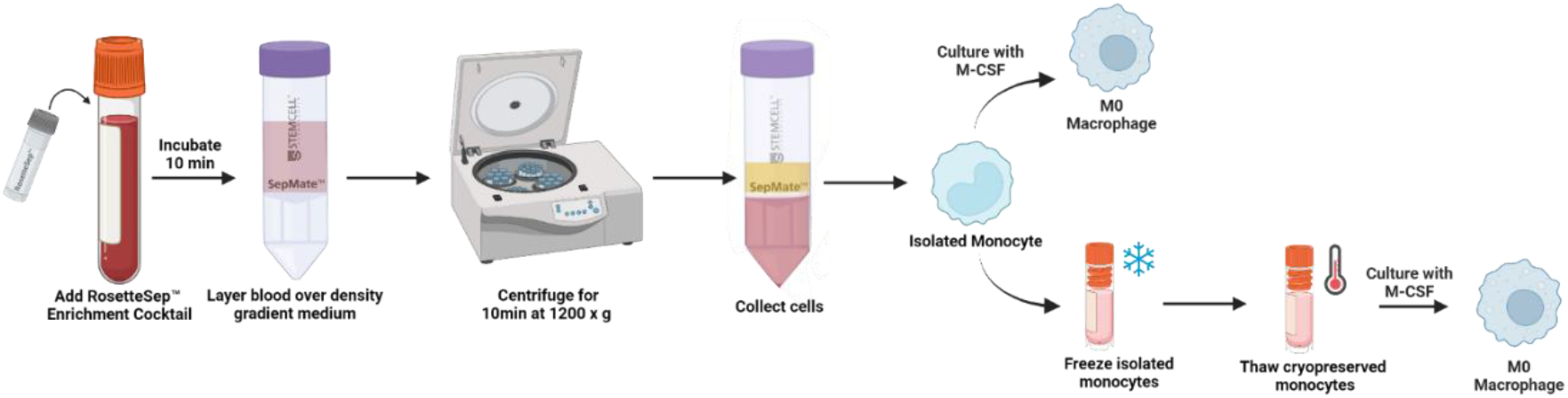
Immune cell isolation using density gradient separation [Created with BioRender.com].

### 2.2 Differentiation of monocytes into macrophages

The isolated monocytes were counted using a standard automatic cell counting method and Trypan blue staining. The monocytes were transferred into 35 mm glass-bottom dishes for culture. The cells were plated at a density of 300,000 cells/dish along with 2 mL of prepared differentiation media, which consists of RPMI −1640 medium supplemented with 10% heat-inactivated FBS and 2 ng/mL human M-CSF. The cells were cultured for 8 days at 37 °C with 5% CO_2_, with media replacement every 2 days following a published differentiation protocol [6].

### 2.3 Cryopreservation and cell thawing cycle

A solution of 5% DMSO in FBS was prepared and kept cold on ice before starting the cryopreservation protocol. After the last centrifugation cycle mentioned in **Methods 2.2**, the supernatant was carefully aspirated without disturbing the cell pellet. The isolated monocytes were then resuspended in 1350 µL of cold FBS and 1350 µL of the 5% DMSO in FBS solution to achieve a 1:1 ratio. This solution was transferred into cryogenic vials that were placed in a −80 °C freezer for several hours before being moved to a cryogenic tank for long-term storage.

The thawing of monocytes followed a standard cell thawing protocol where the cryovials were swirled in a 37°C water bath for 1 minute. The solution in the vials was transferred into bigger centrifuge tubes and 2 mL of monocyte differentiation media was added to the tube dropwise. The samples were centrifuged for 5 minutes at 300 x g. The supernatant was aspirated and the cell pellet obtained was resuspended using 1 mL of differentiation media. The thawed cells were then counted using Trypan Blue staining and an automatic cell counter before being plated in dishes at a density of 300,000 cells/dish.

### 2.4 Imaging

A custom-built multiphoton fluorescence lifetime microscope (Marianas, 3i) was used to excite NAD(P)H and FAD fluorescence at 750nm and 890nm, respectively. Additionally, a 100x objective (NA= 1.46) was used along with 447/60nm and 550/88nm emission filter to isolate these signals. Florescence lifetime images were acquired using a dwell time of 50 µs and 1 frame repeat. Environmental conditions were maintained during imaging experiments using a stage top incubator (Okolab) at 37°C with 85% humidity and 5% CO_2_.

### 2.5 Data analysis

The images captured were analyzed using SPCM software (Becker & Hickl) at the single-cell level. Cell masks were gathered for each NAD(P)H fluorescence image using Cellpose. These images were further processed using MATLAB code to extract meaningful FLIM parameters and compare the metabolic status of fresh and cryopreserved macrophages. The final data was plotted using Rstudio and a two-sided t-test was performed with a p value of 0.05 to evaluate statistical significance.

### 3. Results

Human blood isolated monocytes were differentiated into macrophages through culture with M_CSF cytokine over a period of 8 days. The cells were excited at 750 nm and 890 nm to obtain NAD(P)H and FAD fluorescence data, respectively. Representative images for NAD(P) H and FAD mean lifetime at the 4 imaging time points show a visible increase in the size and changes in the morphology of the macrophages over the 8 days of culture (**Figure 2** and **Figure 3)**.

**Figure 2.**
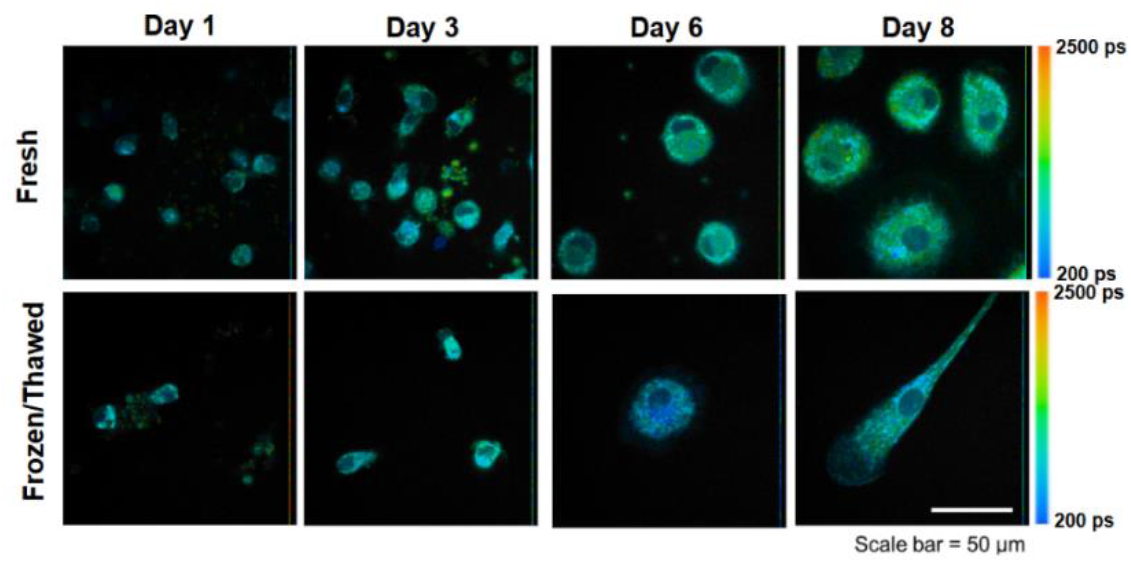
NAD(P)H mean lifetime images of fresh and frozen macrophages captured using multiphoton fluorescence lifetime microscopy.

**Figure 3.**
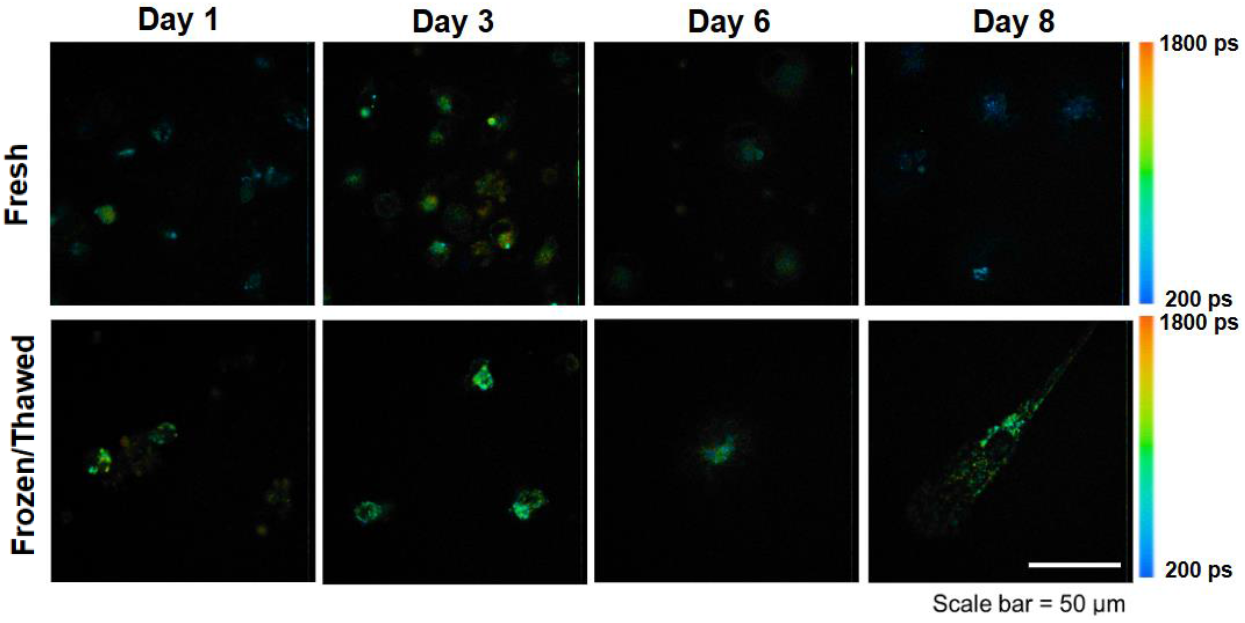
FAD mean lifetime images of fresh and frozen macrophages captured using multiphoton fluorescence lifetime microscopy.

Quantification of the FLIM images revealed a decrease in the mean NADH lifetime between the fresh macrophages and frozen-thawed macrophages at each time point (Fig. 4). In contrast, the mean FAD lifetime increased in frozen-thawed macrophages on day 1 and day 8 as compared with the fresh macrophages (Fig. 4). Similarly, the FAD intensity was increased for the frozen-thawed macrophages as compared with the fresh macrophages at each time point (Fig. 5). The NADH intensity consistently increased over time for both the fresh and frozen-thawed macrophages. The redox ratio significantly decreased at each time point for the frozen-thawed macrophages compared with the fresh macrophages.

**Figure 4.**
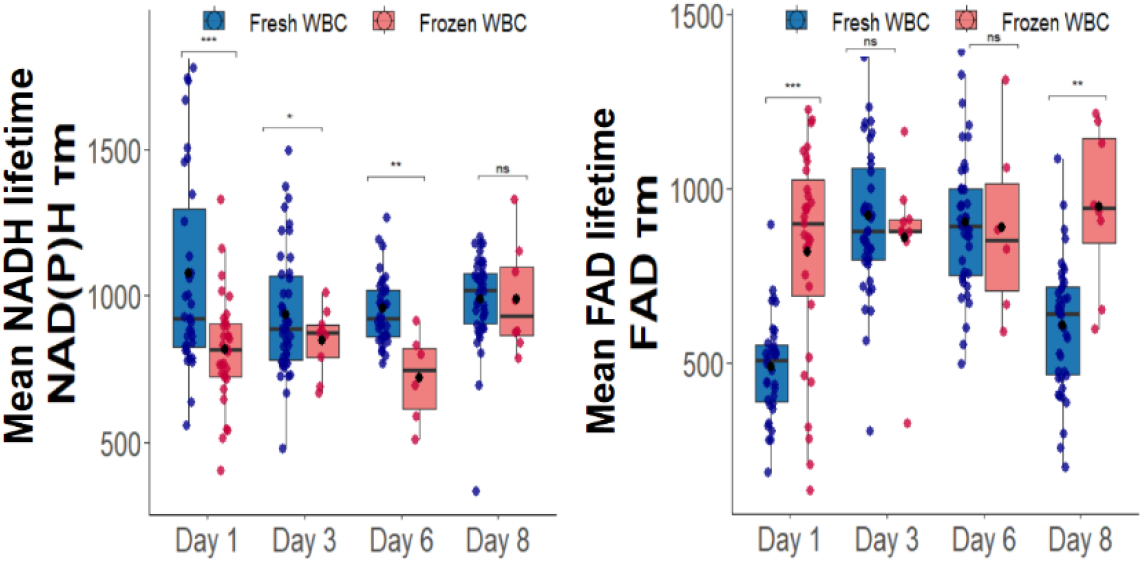
Mean fluorescence lifetime data plots for NADH and FAD of fresh and cryopreserved macrophages.

**Figure 5.**
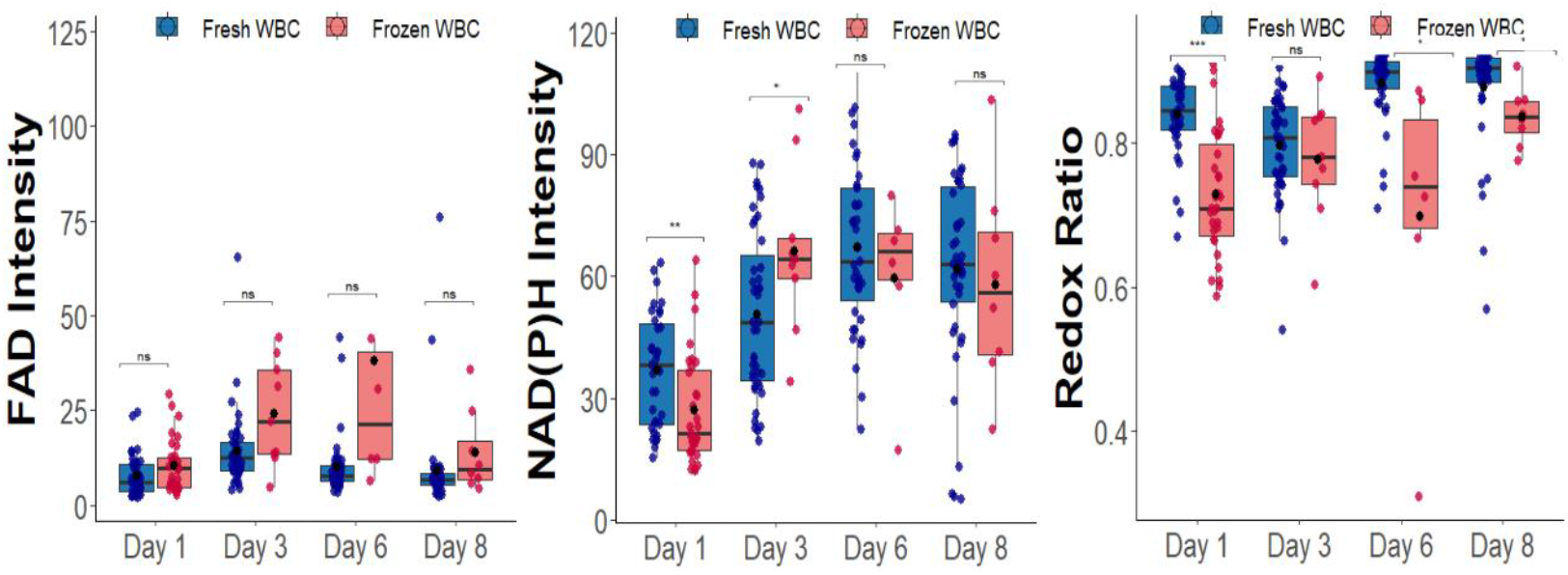
Intensity and redox ratio data plots for NADH and FAD of fresh and cryopreserved macrophages.

## 4. Conclusion

Significant changes in the FLIM parameters between fresh and cryopreserved samples of macrophages suggest that the freezing and thawing cycle of cryopreservation alters the metabolism of macrophages. These changes may be a result of a smaller cell count in the frozen/thawed macrophage group due to a decrease in cell viability from the cryopreservation process.

## Acknowledgements

The funding sources for this research project include Chan Zuckerberg Initiative (2022-251380), NIH-NIGMS (R35 GM142990), and Texas A&M University.

## Notes

### Competing Interest Statement

The authors have declared no competing interest.

### Summary of Updates

This version of the manuscript has been revised to update the abstract.

